# Effective conservation decisions require models designed for purpose: a case study for boreal caribou in Ontario’s Ring of Fire

**DOI:** 10.1101/2022.06.01.494350

**Authors:** Matt Dyson, Sarah Endicott, Craig Simpkins, Julie W. Turner, Stephanie Avery-Gomm, Cheryl A. Johnson, Mathieu Leblond, Eric W. Neilson, Rob Rempel, Philip A. Wiebe, Jennifer L. Baltzer, Josie Hughes, Frances E.C. Stewart

**Affiliations:** Biology Department, Wilfrid Laurier University, Waterloo, ON, Canada; National Wildlife Research Centre, Environment and Climate Change Canada, Ottawa, ON, Canada; University of British Columbia, Vancouver, BC, Canada; Northern Forestry Centre, Canadian Forest Service, Natural Resources Canada, Edmonton, AB, Canada; FERIT Environmental Consulting, Thunder Bay, ON, Canada; Great Lakes Forestry Centre, Canadian Forest Service, Natural Resources Canada, Sault Ste. Marie, ON, Canada

**Author notes:** CORRESPONDENCE: Frances Stewart, Wilfrid Laurier University, Biology Department 75 University Ave., Waterloo, Ontario, N2L 3C5. shared senior authorship as Co-PIs (Josie Hughes, Frances E.C. Stewart).

**Keywords:** *Rangifer tarandus caribou*, Open science, Decision-support, Resource selection function, Demographic model, environmental impact assessment, caribouMetrics

## Abstract

Decision making in conservation science often relies on the best available information. This may include using models that were not designed for purpose and are not accompanied by an assessment of limitations. To begin addressing these issues, we sought to reproduce, and evaluate the suitability of, the best available models for predicting impacts of proposed mining on boreal woodland caribou (*Rangifer tarandus caribou)* resource selection and demography in northern Ontario. We then evaluated their suitability for projecting the impacts of development in the Ring of Fire region. To aid in accessibility, we developed an R package for data preparation, analyses of resource selection, and demographic parameters. We found existing models were either ill suited, or lacking, for ongoing regional planning. The specificity of the regional resource selection model limited its usefulness for predicting impacts of development, and the high variability across caribou ranges limited the usefulness of a national aspatial demographic model for predicting range-specific impacts. Variability in model coefficients across caribou ranges suggests selection responses vary with habitat availability (i.e. a functional response) while demographic responses continue to decline with increasing disturbance. Models designed for forecasting that are continuously updated by range-specific demographic and habitat information, are required to better inform conservation decisions and ongoing policy and planning practices in the Ring of Fire region.

## 1 INTRODUCTION

Decisions about natural resource development should be informed by the best available information (Fuller et al. 2020). Models that quantitatively link wildlife species responses to development, such as wildlife-habitat relationships and demography (Beyer et al., 2010; Matthiopoulos et al., 2020), can inform these decisions and associated policies (Wilson et al., 2021). However, these tools are generally not designed for decision support, and decision makers may be unaware of their limitations. For example, models are rarely reproducible; data and modelling processes are rarely reported nor accessible without author engagement, resulting in limited applicability outside of the specific context for which models were built. These problems can be addressed by couching models within an open source, and easily reproduced, decision-support tool (e.g., Eacker et al., 2019; Nagy-Reis et al., 2020; Nowak et al., 2018; McIntire et al. 2022) that can inform impact assessment, particularly for Species at Risk (Roche et al., 2020).

As a country supporting some of the largest intact landscapes globally, Canada has a heightened responsibility to conduct natural resource development decisions in the most sustainable way possible. For example, the Hudson Bay Lowlands of Northern Ontario is one of the largest intact wetlands and peatland carbon stores on the planet (Ibisch et al., 2016; Poley et al., 2022; Sothe et al., 2022; Tootchi et al., 2019). The region is home to many wildlife species and is largely inaccessible by road. At the same time, it is rich in valuable minerals and is a target for resource extraction and economic development aimed at a green energy transition (Carlson and Chetkiewicz, 2013; Far North Science Advisory Panel (Ont.) et al., 2011; IAAC, 2021). With an anticipated Regional Assessment in the region (IAAC, 2023), and much interest in critical minerals for batteries and other purposes (NRCan 2022), there is a need for modelling tools to assess the potential environmental impacts of the proposed mining of ‘Ring of Fire’ (RoF) mineral deposits and associated development on wildlife in this globally significant ecosystem (Far North Science Advisory Panel (Ont.) et al., 2011; IAAC, 2021; Fig. 1).

**Figure 1.**
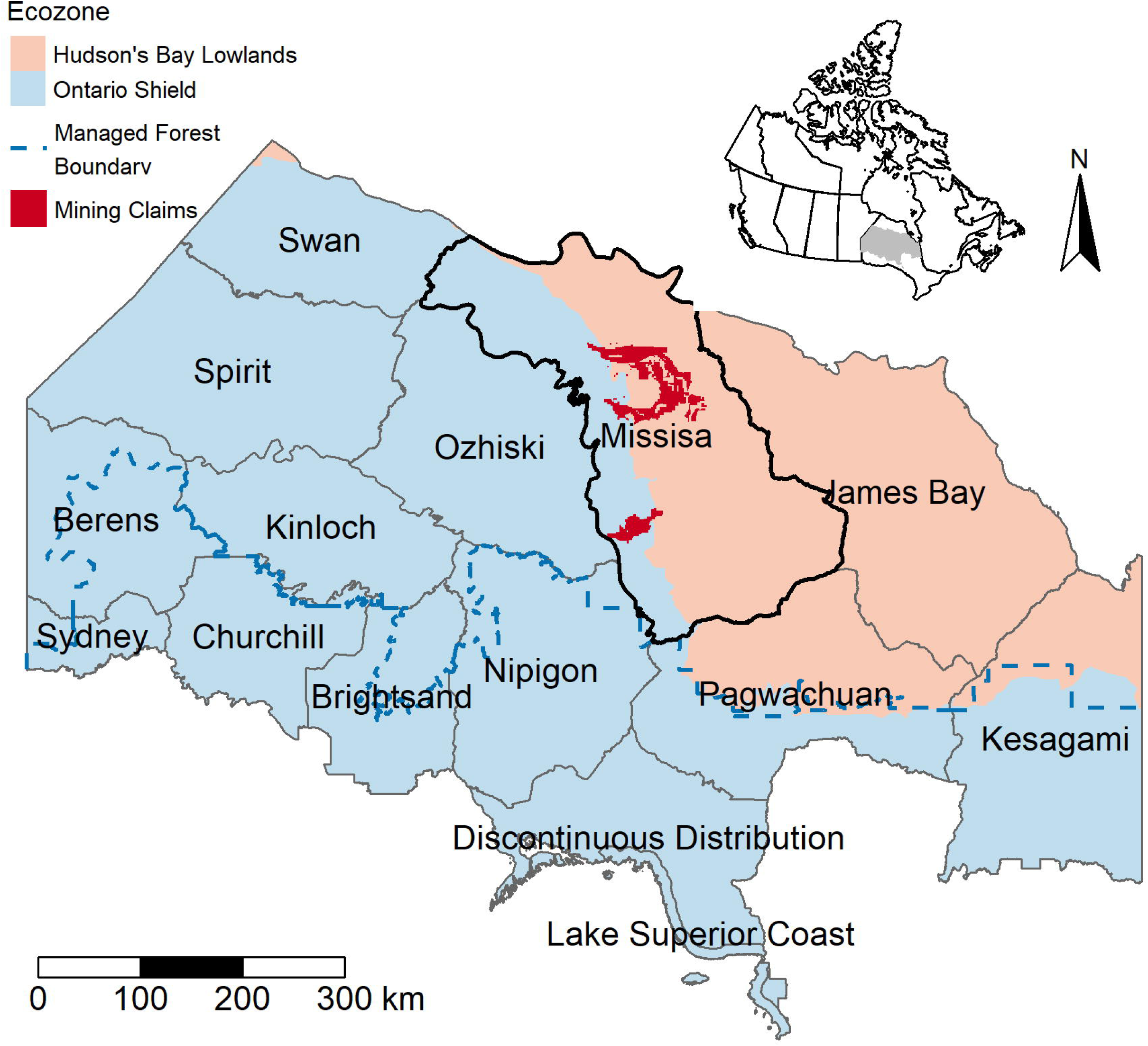
The Missisa boreal caribou range (black outline), which includes the proposed Ring of Fire (RoF) mining claims (dark red), is the focus of this study. The blue dashed line distinguishes the Managed Forest Zone (MFZ) from the Far North, and shading distinguishes the Ontario Shield ecozone (pale red) from the Hudson Bay Lowlands(blue). Inlay map (top-right) indicates the location of the Ontario boreal caribou ranges (grey) relative to Canada.

Quantitative knowledge of wildlife-habitat relationships are commonly represented by resource selection functions (RSFs), which can produce maps of habitat suitability through space and time, and can infer consequences of habitat loss (Boyce et al., 2002; Harju et al., 2011; Johnson et al., 2004; Matthiopoulos et al., 2019). However, RSFs represent descriptive associations between the behaviour of a sample of animals and habitat under current, scale-specific conditions, limiting their predictive capacity and transferability to novel conditions (Boyce, 2006; Decesare et al., 2012b; Kunegel-Lion et al., 2022; Yates et al., 2018). Aspatial wildlife demographic models are another tool and provide insight into drivers of population growth, such as survival and recruitment, and are also used for characterizing demographic responses to landscape alteration (Johnson et al., 2020; Sorensen et al., 2008; Stewart et al., 2020). However, high uncertainty about demographic parameters, drivers of change, and vital rates can limit the utility of models for decision making and regional application (Chaudhary and Oli, 2020; Sleep and Loehle, 2010; but see Stewart et al. 2020).

Boreal populations of woodland caribou (*Rangifer tarandus caribou*; hereafter ‘boreal caribou’) inhabit large portions of Canada’s boreal forest (ECCC, 2011, 2019; Festa-Bianchet et al., 2011), where anthropogenic disturbance from natural resource extraction sector threatens population persistence (Fryxell et al., 2020; Hebblewhite, 2017; Stewart et al. 2020; Johnson et al., 2020). Nationally, only 29% of population ranges (15 of 51) are considered self-sustaining and boreal caribou are listed as threatened under the federal Species at Risk Act (ECCC, 2024a). In the province of Ontario, boreal caribou are listed as threatened due to development from forestry and mineral exploration, including their road networks, which together increase predation and reduces adult female and calf survival (MNRF, 2014a) Ongoing landscape disturbances are predicted to cause continued population declines (Fryxell et al., 2020). Effective conservation of caribou in Ontario requires spatial and demographic tools that can project the effects of future landscape and climatic changes on caribou habitats and population performance. An existing Ontario RSF (Hornseth and Rempel, 2016) and national demographic model (Johnson et al. 2020) represent the best available regional prediction tools for this species but have many potential shortcomings (ECCC 2024b): notably, they rely on a limited amount of data from over a decade ago, and their utility for transferability to other contexts and projecting anticipated anthropogenic impacts has not been evaluated.

To advance the practice of conservation science decision making for a species at risk in a changing landscape, we sought to evaluate the suitability of the best available models for predicting impacts of proposed mining in the RoF on Ontario’s northern boreal caribou. To assess the potential impacts, we reproduced existing caribou RSF and demographic models and evaluated their suitability for spatial transferability and prediction. Species at risk models can be difficult to fully reproduce as information is rarely made publicly available. In this context, location data for the RSF was not available and error estimates for model coefficients were not published (Hornseth and Rempel 2016). For the national demographic model, code and regression model parameters were made available upon request from the authors (Johnson et al., 2020), but data owned by provinces was not accessible. We demonstrate and discuss the challenges of this exercise and suggest open source tools and reproducible workflows as a solution by reproducing the existing models within an R package framework (Wickham, 2015). To aid this effort, we created an R package (caribouMetrics) that can be integrated into predictive frameworks to support resource use decisions or modified and advanced by other practitioners for other uses (Bodner et al., 2021a; McIntire et al., 2022; Micheletti et al., 2021; Miller and Frid, 2022).

## 2 METHODS

### 2.1 Study Area

We focused on four caribou ranges, as delineated by Ontario’s provincial government, that are adjacent to or overlap with the RoF mining claims and had published RSF coefficients: Missisa, James Bay, Pagwachuan, and Nipigon. The Missisa range contains most of the RoF mineral claims (Far North Science Advisory Panel (Ont.) et al., 2011; IAAC, 2021; Fig. 1) and is the focus of our study. Missisa and James Bay ranges occur in Ontario’s Far North (Far North Science Advisory Panel (Ont.) et al., 2011), where anthropogenic disturbance has so far been limited to small communities and exploration camps, winter roads, and the now inactive Victor Diamond Mine. Nipigon and Pagwachuan ranges are almost entirely within the Managed Forest Zone (MFZ) where industrial forestry is common (Fig. 1).

These four ranges straddle the Ontario Shield and Hudson Bay Lowlands ecozones (Fig 1). The Shield consists of mixed and coniferous forests dominated by black spruce (*Picea mariana*), and containing jack pine (*Pinus banksiana*), balsam fir (*Abies balsamea*), white spruce (*Picea glauca*), trembling aspen (*Populus tremuloides*), and balsam poplar (*Populus balsamifera*; Crins et al., 2009). The Missisa and Pagwachuan ranges include the transition between the Shield and Lowlands, which is mainly composed of peatland complexes with poor drainage, and trees including stunted black spruce and tamarack (*Larix laricina*; Crins et al., 2009). The James Bay range is almost entirely within the Lowlands and the Nipigon range is entirely within the Shield (Fig 1).

### 2.2 RSF Model Reproduction & Validation

To reproduce existing RSFs for caribou in northern Ontario we used tables of published boreal caribou RSF coefficients for the same areas as Hornseth and Rempel (2016) (hereafter referred to as the ‘original RSFs’). Hornseth and Rempel referred to these models as resource selection probability functions (RSPFs), but their approach differs from what is commonly understood as an RSPF (Johnson et al., 2006); as such we use the term ‘RSF’ throughout. The data used to produce the original RSFs were collected between 2009 and 2013 from GPS-collared caribou across a variety of companion studies within the ranges of interest; however, these data are not publicly available (Hornseth and Rempel, 2016; MNRF, 2014b, 2014b; Rempel and Hornseth, 2018). Instead, we obtained the published coefficients for range-specific seasonal RSFs of each reported top model (Table S1.1) for the Nipigon, Pagwachuan, Missisa, and James Bay ranges. However, associated error or uncertainty information for individual coefficients was not reported in Hornseth and Rempel (2016) and model performance was only reported at the model level (e.g., sensitivity, specificity, and area under the receiver operating [ ROC] characteristic curve).

The predictors used in the original RSFs were developed from land cover, fire and harvest disturbance history, linear features, and esker datasets (Tables S1.2, S1.3). To reproduce the original RSFs, we acquired data sets for the predictor variables that were publicly available and matched the data sources described by Hornseth and Rempel (2016) as closely as possible. Harvest, fire, and linear features data from the present (2020) were available and we used information on the timing of the event or the feature’s construction to recreate the data that was available when the original models were produced (Table S1.2). We used current versions of these same datasets to project the RSFs under different development scenarios (Table S1.2).

To assess our ability to reproduce the original models, we acquired the original projected surfaces for comparison from the authors (R. Rempel pers. comm, 2021). We validated our models visually and quantitatively. The original models used a nested hexagon grid, common in Ontario (e.g. Poley et al. 2014; Ray et al. 2018), to generate predictions. For ease of future implementation using raster tools, we approximated this approach with distance-weighted moving windows on a rectangular grid, and compared the two approaches by extracting the values from our rectangular grid to the 592-hectare hexagonal grid provided by the authors using the weighted mean. We mapped the difference in predictions and explanatory variables between the models and used scatterplots to compare the values of the model responses for each grid cell produced by both methods. A perfect reproduction would produce a Pearson correlation coefficient of 1, and any deviation from the original prediction would reduce this value. We expected some differences in the predictions to result from the type of grid and predictor data sets used.

### 2.3 Disturbance Scenarios and RSF Model Projection

To assess the suitability of existing RSFs for projecting the impacts of disturbance on boreal caribou habitat selection we considered three simple disturbance scenarios. We created a ‘base’ scenario with no change in disturbance footprint (0.41% anthropogenic disturbance), a ‘roads-only’ scenario that included proposed RoF access roads (1.11 % anthropogenic disturbance; MNDMNRF, 2022), and a ‘roads-and-mines’ scenario that included the proposed access roads and mining claims associated with the RoF (6.95% anthropogenic disturbance; Fig. 2; Table S1.2). The original RSF included road density as a predictor and required additional information on roads within mining areas that we were not able to obtain; we therefore only compared the base and road-only scenarios for the RSF but included all three scenarios for the demographic model (see below).

**Figure 2.**
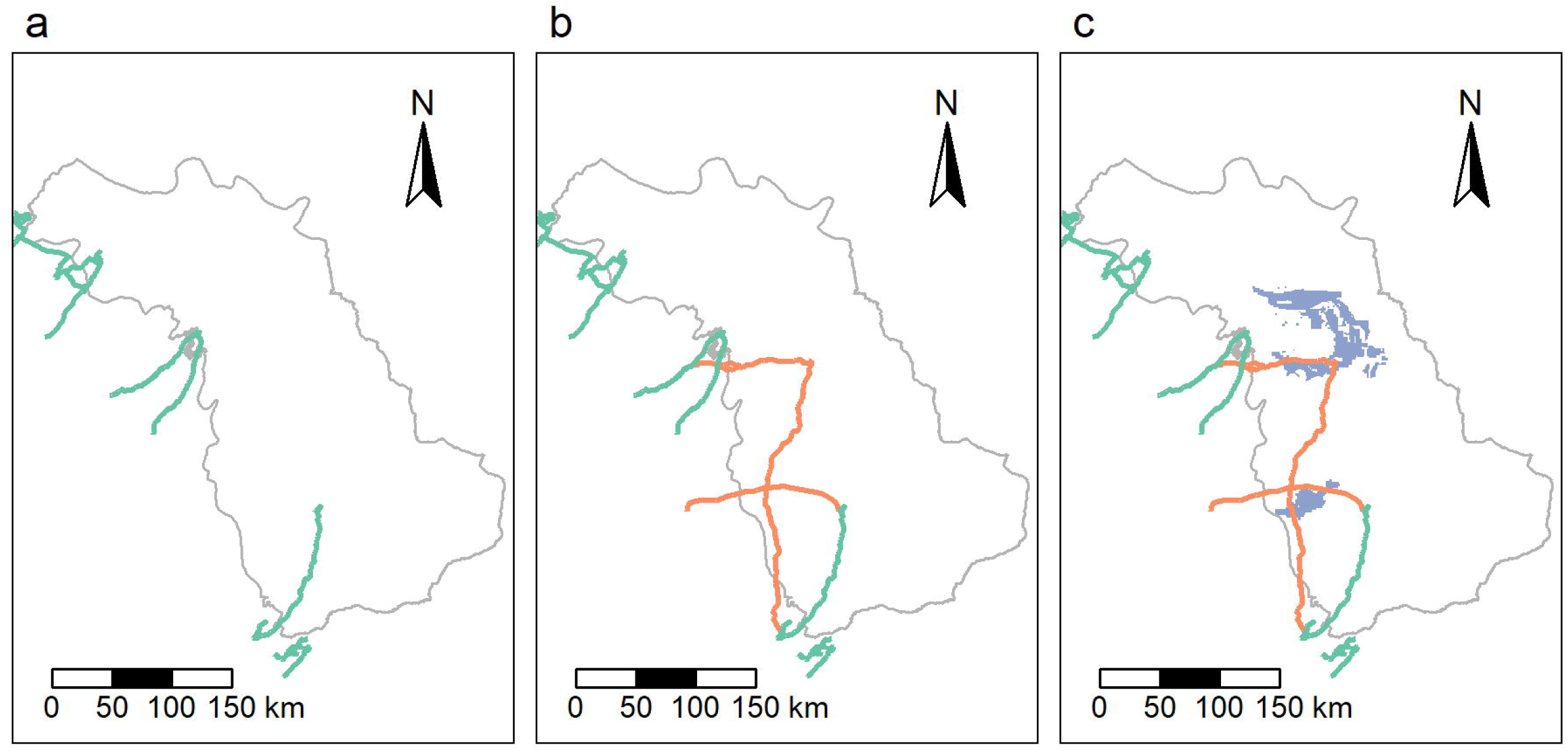
Maps of the Missisa range with the extent of linear features included in (a) the original model developed by Hornseth and Rempel (base), (b) the roads-only scenario used to project the RSF model, and (c) the roads-and-mines scenario used for demographic modelling. Existing roads are represented as green lines, proposed roads as orange lines, and mines are coloured in purple.

To understand how the ‘roads-only’ scenario might affect resource selection, we projected the original RSFs across updated landscape conditions as represented by i) temporal changes in forest structure between 2010 and 2020 (e.g., including simulated fires; Table S1.2) and ii) the new proposed roads-only scenario (Fig. 2). We also assessed the potential for borrowing RSF information from other ranges by transferring coefficients from the top original RSFs from adjacent ranges (James Bay, Nipigon, and Pagwachuan) to Missisa. We visualized the spatial transferability of these models, and then examined their sensitivity to changing habitat availability using scatterplots where we compared the value of the model response for each grid cell produced by the Missisa model and each respective adjacent range.

### 2.4 Demographic Model Reproduction, Validation, and Projection

Canada’s national demographic boreal caribou model was developed from adult female survival (*S*), calf recruitment (*R*), and landscape data across 58 boreal caribou study areas, including 13 study areas in Ontario (Johnson et al., 2020; Table S1). We used models (Johnson et al., 2020) to predict changes in *S* and *R* within the roads-only and roads-and-mines disturbance scenarios. The national demographic model is aspatial and all types of anthropogenic disturbances are combined into a single measure of disturbed area within a range; using the ‘roads-and-mines’ scenario is sufficient and there is no need to specify the location of roads within mining claims. We calculated the relevant predictor variables for the Missisa range based on Johnson et al. (2020;i.e., % anthropogenic disturbance buffered by 500 m; % wildfire within the last 40 years; Table 1), and calculated expected *R* and *S* as a function of disturbance according to the beta regression models with highest support (M4 and M1 respectively from Johnson et al., 2020):

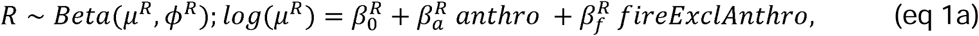

**Table 1.** Summary of changes in buffered anthropogenic disturbance, and fire excluding anthropogenic disturbance (as defined in Johnson et al. 2020), in three scenarios: the Base Scenario without additional development, Roads Only scenario, and Roads-and-Mines scenario that included the proposed roads and mining claims within the Ring of Fire area.

| Scenario | Anthropogenic (%) | Fire (%) | Total Disturbance (%) | Fire Excluding Anthropogenic (%) |
| --- | --- | --- | --- | --- |
| Base | 0.41 | 4.27 | 4.58 | 4.17 |
| Roads Only | 1.11 | 4.27 | 5.22 | 4.12 |
| Roads-and-Mines | 6.95 | 4.27 | 11.02 | 4.71 |

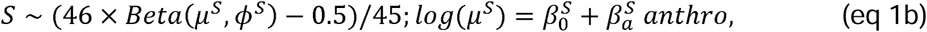

Where □*^R^ ∼ Normal(19.862,2.229)* and □*^S^ ∼ Normal(63.733,8.311)* are precisions of the Beta distributed errors (Ferrari and Cribari-Neto, 2004), and survival rates are back transformed as in Johnson et al. 2020. Table 3 of Johnson et al. (2020) provides the expected values and 95% confidence intervals of all regression coefficients (β*_0_^R^,* β*_α_^R^,* β*_f_ ^R^,* β*_0_^S^,* β*_α_^S^*), which are assumed to be Gaussian distributed. To evaluate these recruitment (eq 1a) and survival (eq 1b) models, we sampled expected demographic rates across a range of anthropogenic and fire disturbance (0-100%) to reproduce expected values and 95% predictive intervals from Fig. 3 and Fig. 5 of Johnson et al. (2020).

**Figure 3.**
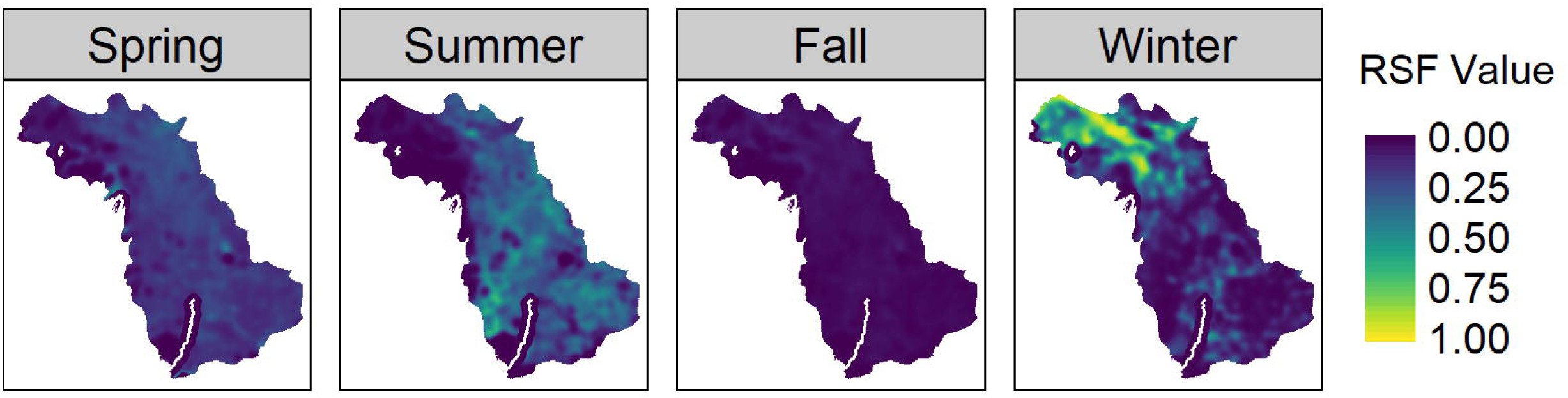
Reproduction of the seasonal RSF for the Missisa range from Hornseth and Rempel (2016) using *caribouMetrics* and the published coefficients to reproduce the relative probability of use (0-1) by boreal caribou during spring, summer, fall, and winter. The predictor variables used are approximations of those used by Hornseth and Rempel (2016) based on currently available data. Scale ranges from dark blue to yellow with yellow representing a higher relative probability of use.

In areas of low anthropogenic disturbance, such as the Missisa and James Bay ranges, there is substantial among-population variability in recruitment (Fig. 3 in Johnson et al., 2020). To model this variation, we selected regression model parameter values for each study area (i.e. a sample population) at the beginning of simulations and assigned to quantiles of the error distributions for survival and recruitment. Sample populations remained in their quantiles as the landscape changed, allowing us to distinguish the effects of changing disturbance from variation in initial population status. To show these effects, we projected population growth for 35 sample populations across a wide range of anthropogenic disturbance levels (0-90%). We slightly altered Johnson et al’s (2020) demographic model by including demographic stochasticity (Supplementary Materials Part 2; see Hughes et al. *in review* for details).

We characterized the distribution of outcomes for our three disturbance scenarios (Fig. 2, Table 1) more thoroughly by projecting the dynamics of 500 sample populations. Population growth rate each year is given by □*_t_=N_t+1_/N_t_* when *N_t_>0* and □*_t_=0* when *N_t_=0*. For each sample population, population growth was projected for 20 years from an assumed initial population size of 373 females, which we derived from the minimum animal count of 745 caribou between 2009 and 2011, assuming 50% were female (MNRF, 2014b). This estimate is conservative because adult male:female ratios are typically lower in caribou (ECCC, 2011; Espinoza and Weckerly, 2021). We report the average growth rate., over that time. To verify our reproduction of the demographic model used by Johnson et al. (2020), we compared our outputs to those from model code supplied by the authors.

### 2.5 caribouMetrics: an R Package for Model Reproduction, Validation, and Projection

We incorporated RSF and demographic model components into the *caribouMetrics* R package with documentation and vignettes explaining their use and used GitHub to promote version control and transparency of the development process. This package can be accessed on GitHub: https://github.com/LandSciTech/caribouMetrics.

The standardized nature of packages allows them to easily integrate into the larger R ecosystem, facilitating ease of extension and adaptation for tasks beyond their original design (e.g. prediction or forecasting; McIntire et al., 2022). *caribouMetrics* also contains functions that automate the geospatial data preparation process to facilitate application to new landscapes. The original RSFs were developed using a closed scripting language called Landscape Scripting Language (LSL; Kushneriuk and Rempel, 2011), which we re-implemented in R. For the demographic model, we began by borrowing demographic rate sampling code from a SpaDES (Chubaty and McIntire, 2022) module (*caribouPopGrowth*; https://github.com/tati-micheletti/caribouPopGrowthModel; Stewart et al., 2023), and advanced the method to include precision and quantiles. Together, this makes the national caribou demographic model (Johnson et al. 2020) more transparent, reproducible, and adaptable to local population estimates.

## 3 RESULTS

### 3.1 RSF Model Reproduction, Validation, and Projection

We were able to produce a reasonably accurate representation of the original RSFs for the Missisa range, as evidenced by high Pearson’s r values across all seasons (r > 0.935; Fig. S1.1). However, some locations showed different predictions depending on the season (Fig. S1.2). This was expected given the input data was not exactly the same between the two methods (Fig. S1.3). In the Missisa range, the model predicted the highest relative use probabilities in the northwest of the study area during winter (Fig. 3), consistent with the original RSFs. During the summer and spring, the eastern portion of the range was used more than the northwest (Fig. 3).

We observed a lower relative probability of use in areas associated with proposed roads (Fig. 4). Lower relative probability of use along the road corridors was strongest in the spring and summer, consistent with seasonal changes in the response to roads described in the original RSFs (Fig. 4). There was high variability in the estimated response to roads among ranges (Fig. 4). The James Bay range prediction appeared the most similar to the Missisa range prediction (Fig. 4); however it varied by season (Fig S1.4). The Nipigon and Pagwachuan range projections, which visually differed substantially from the Missisa projection, did not show a strong response to proposed roads (Fig. 4; Fig S1.4).

**Figure 4.**
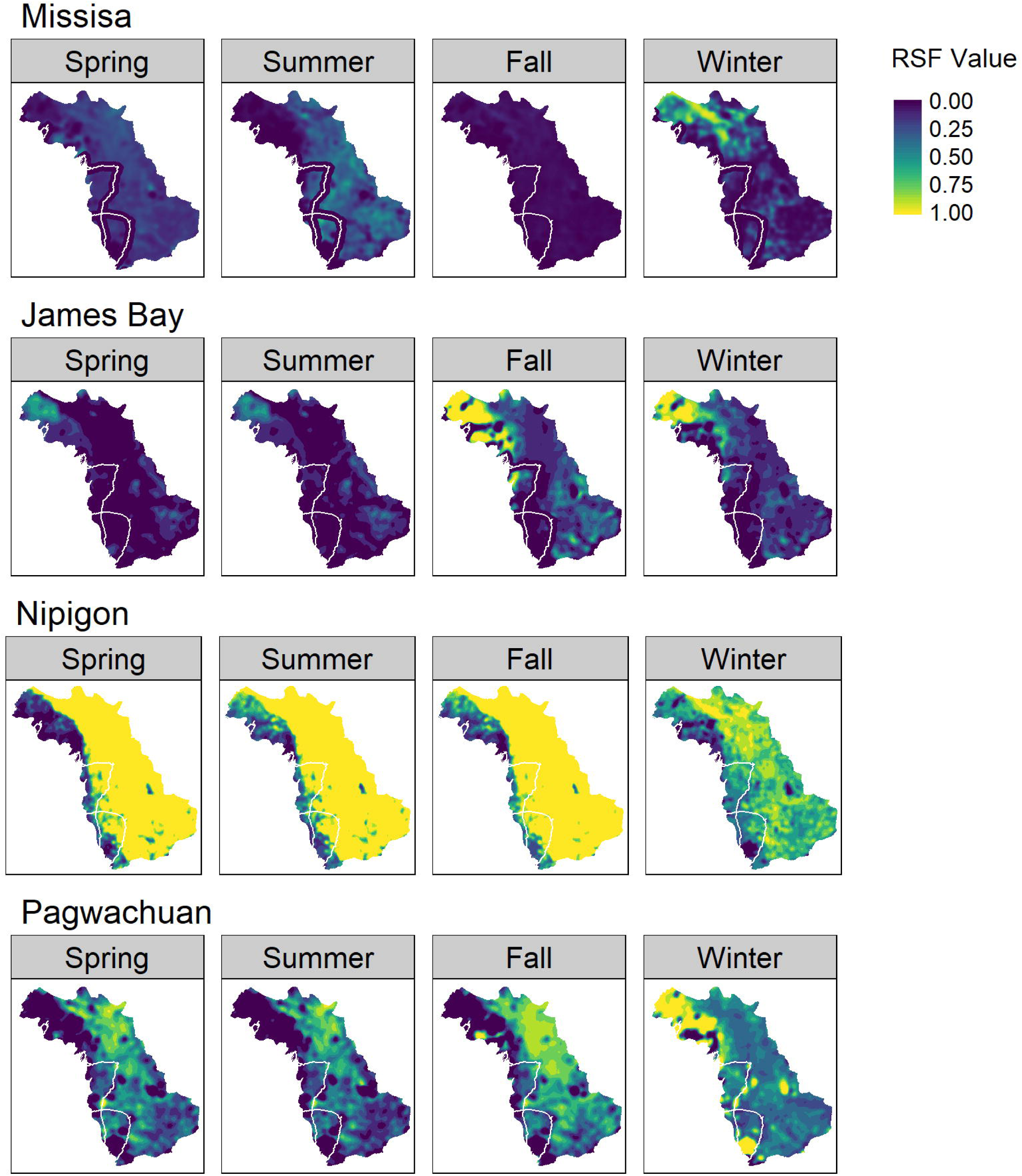
Seasonal RSF predictions from *caribouMetrics* in the Missisa range under the roads only scenario using the coefficients from Hornseth and Rempel (2016) from the Missisa, James Bay, Nipigon, and Pagwachuan range to estimate the relative probability of use (0-1) by boreal caribou during spring, summer, fall, and winter. Scale ranges from dark blue to yellow with yellow representing a higher relative probability of use.

### 3.2 Demographic Model Reproduction, Validation, and Projection

Our regression models for survival and recruitment closely matched those of Johnson et al. 2020 (Fig. 5). Anthropogenic disturbance remained low in all our disturbance scenarios (Table 1), and the corresponding range of variability in demographic rates among sample populations was high (Fig. 5). The model predicted that increasing anthropogenic disturbance would decrease both survival and recruitment (Fig. 5), but the importance of that decrease for self-sustainability of the population was highly uncertain and depended on initial population status (Fig. 5).

**Figure 5.**
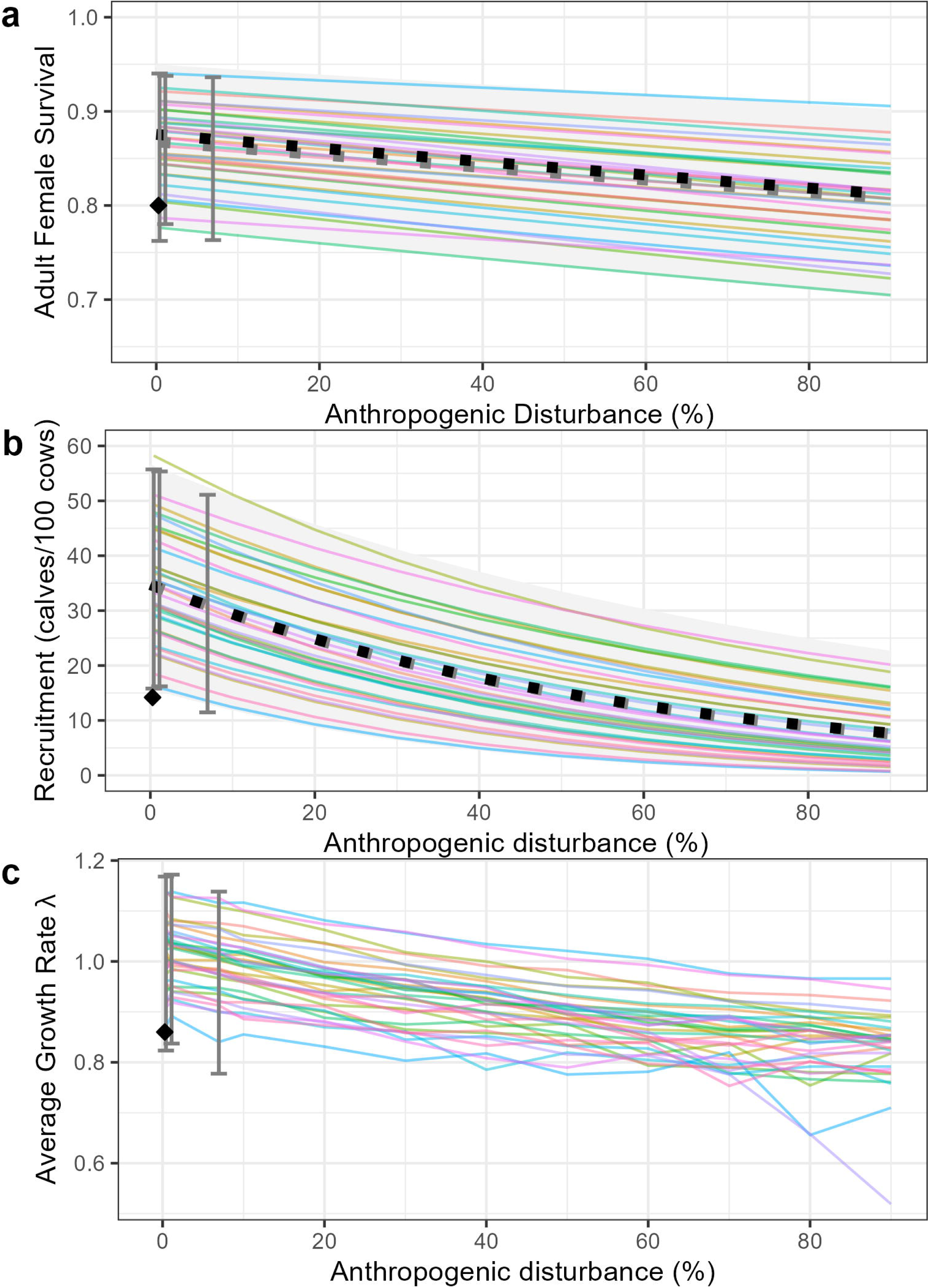
Demographic rate simulations derived from regression models in Johnson et al. (2020) for (a) Adult female survival (S), (b) Recruitment (calves per 100 cows), and (c) Average population rate (.,). Overlap of grey and black dotted lines in (a) and (b) indicates a good match between expected values from our model (grey) and Johnson et al. 2020 (black). Coloured lines show effects of changing anthropogenic disturbance on demographic rates in 35 sample populations, assuming sample populations are randomly distributed among quantiles of the beta distribution, and each population remains in the same quantile of the beta distribution as disturbance changes. Alignment of these sample trajectories with 95% predictive intervals from Johnson et al. 2020 (pale grey bands in panels a and b) indicates that we have adequately reproduced variability in that model. Bars show the 2.5th and 97.5th percentiles for 500 sample populations under the three disturbance scenarios for the Missisa range (from left to right, Base, Road Only, and Roads and Mines). The black diamond indicates the demographic rates for the Missisa range according to a 2014 assessment (MNRF, 2014a). Populations with an average population trend of less than 0.99 are considered not self-sustaining.

## 4 DISCUSSION

To evaluate existing models and assess their limitations for projecting anticipated landscape impacts we examined the applicability of two kinds of existing models; resource selection functions (RSF) and demographic models, for boreal caribou. We found that the original Missisa range RSF was poorly suited for projecting the impacts of disturbance because it was specific to conditions at the time of model development in an area that contained limited anthropogenic disturbance. This led to high uncertainty in the estimated impacts of linear features, with a low standardized effect size, high variability among seasons, and the possibility that the model understinated future impacts of linear features. In contrast, the existing national demographic model was too general to project the impacts of disturbance in this range. Additional data collection is necessary but not sufficient to address these limitations (see also ECCC 2024b); there is also a need for new models that are better suited to forecasting the impacts of disturbance. These models should be designed for purpose. They should identify and account for uncertainty, be updateable with new information (e.g., local data), be transparent, simple, and reproducible (Bodner et al., 2021b).

### 4.1 Lessons learnt from using non-predictive models to forecast impacts

Across caribou ranges where anthropogenic disturbance is low, the national demographic model predicted high variation in recruitment and survival among populations (Fig. 3 and Fig. 5 of Johnson et al., 2020), leading to high variability in projected impact of increasing disturbance on population growth rate in the RoF (Fig. 5 here). This variability highlights the need for local information about the current (i.e., baseline) demographic parameters of caribou populations in the RoF and ongoing monitoring of disturbance impacts in the area. An assessment of the status of the Missisa population based on 2008 – 2012 data (MNRF, 2014b) suggested survival, reproduction, and population growth rate in this range were lower than predicted by the national demographic model of Johnson et al. 2020 (see black diamond in Fig. 5). The national demographic model (Johnson et al., 2020) predicts that increasing anthropogenic disturbance will decrease both survival and recruitment, but the importance of that decrease for population viability depends on baseline demographic parameters. Even small changes in adult female survival can affect the sustainability of a population in ranges with low adult survival (Johnson et al., 2020). We a method for integrating local survival, recruitment, and status information into range-specific demographic projections informed by national demographic-disturbance relationships, but see Hughes et al. *in review* for recent advancements. Without adequate recent information on population size and demographic parameters, it is difficult to predict the impacts of proposed RoF disturbance on the viability of this particular population with any degree of confidence.

### 4.2 Modelling challenges and opportunities

The original RSF models (Hornseth and Rempel, 2016) were fit independently with data from each range. This is a reasonable approach when the objective is to characterize current habitat use, but not for projecting responses to changing landscape conditions. In the original RSFs there was high variability in the effect of linear features among range-specific RSFs, suggesting that the behavioural response of caribou to linear features may vary with the amount or type of disturbance in a range (i.e., a functional response; Mysterud and Ims, 1998), or that effects of linear features were confounded with other correlated predictors (e.g., forest harvest; Hornseth and Rempel, 2016). Hierarchical regional models that include functional responses to disturbance (Matthiopoulos et al., 2011; Muff et al., 2020; Olson et al., 2021; Teitelbaum et al., 2021) could yield models that are better suited for projection.

Our method of modelling variation in survival and recruitment among populations (Fig. 5) adequately reproduced the observed variation among populations (Fig. 3 and Fig. 5 in Johnson et al., 2020), though it differs from the range-specific scenario approach taken by Johnson et al. (2020). We assumed demographic parameters (i.e., recruitment and survival) vary with disturbance according to the best supported national models, but we note that other competing models with comparable support (i.e., Table 2 of Johnson et al., 2020) are also plausible and might yield somewhat different projections when applied to northern ranges with low anthropogenic disturbance (*sensu* Stewart et al. 2023). This simple population model assumes no variation in recruitment or survival with age, sex, or other parameters (Supplementary Material Part 2). The simple demographic model also ignores uncertainty associated with imperfect survey methods (DeCesare et al., 2012a; Ellington et al., 2020). A more thorough investigation of the sensitivity of caribou demographic projections to variation in model form and measurement errors is warranted but beyond the scope of this paper.

### 4.3 Next steps for supporting actionable science in the Ring of Fire region

In the Ring of Fire region, existing data and models are not sufficient for forecasting changes in caribou habitat selection and demography. Recent caribou movement and demographic data is lacking, and there are also important deficits in our understanding of disturbance history, impacts, and recovery in the low productivity peatlands of the Hudson Bay Lowlands. The region is incredibly remote, making data collection expensive logistically difficult. There are various efforts underway to collect data needed to inform resource development and wildlife management decisions in the RoF (e.g., Government of Canada, 2022), and we hope these efforts also lead to opportunities for collaborative development of region-specific open-source forecasting models. We hope our analysis of existing models helps clarify that new data and models are needed to inform decisions in this region; existing models are not enough to provide spatially explicit information to minimize detrimental effects of human development on caribou recovery in northern Ontario

Although we believe that more data is needed in the Ring of Fire region, it does not follow that more data is needed in all regions and circumstances. We advocate for an open, transparent, iterative modelling approach that allows for integration of data and models as they become available, *sensu* caribou metrics and Hughes et al. (*in review*). This approach allows ongoing assessment of the implications of additional data for model uncertainty and can address important questions about what additional data is required to inform conservation decisions. A well-designed, open, transparent, iterative modelling framework can help support and inform important, nuanced, and region-specific decision making about exactly what data is needed and why.

## 5 CONCLUSIONS

We reiterate a call for improvements in future models, and their workflows, used for conservation decision-making. To assess disturbance impacts on wildlife, models need to be designed for purpose, identify and account for uncertainty, be updateable as new predictor information is available (e.g., local data), and be reproducible (Bodner et al., 2021b). We encourage developers of wildlife response models and collectors of relevant wildlife data to work together toward this goal (Davidson et al., 2021; Russell et al., 2021).

Developing usable decision-support tools demands time and specialized skills that may not be accessible to everyone engaged in open conservation science. One solution is to work in multi-disciplinary teams (Bodner et al., 2021b). Another is to recognize that steps and training toward transparency and reproducibility are valuable, even if the result is not always an easily usable tool. In this project, we were able to reproduce existing models only because the developers of those models were willing to share their code and discuss their model generation procedures. Code does not have to be flawless to enable others to build on previous work, and a shift to more open workflows will reduce the chance of errors, increase efficiency, and advance our ability to generalize across studies and systems (Alston and Rick, 2021; Lewis et al., 2018). We hope that improving the transparency, reproducibility, and decision-relevance of wildlife response models will strengthen our collective ability to make informed decisions and advance conservation science and practice.

## AUTHOR CONTRIBUTIONS

MD led the writing and co-led the analysis; SE and CS co-led package development and provided support for analysis and writing; JT provided contributions to writing and evaluation of the RSF; SAG, CAJ, ML, EN, RR, PW (listed in alphabetical order) provided advice on package development, analysis, and writing; RR and CAJ also provided code and insight into the RSFs and demographic models. FS, JB, and JH provided supervision and oversaw the writing and analysis of this piece.

## DATA AVAILABILITY STATEMENT

The caribouMetrics R package is available at https://github.com/LandSciTech/caribouMetrics, and analysis code is available at https://github.com/LandSciTech/MissisaBooPaper. Data required for analysis is available in this OSF repository: https://osf.io/r9mkp/?view_only=fb71321265d14dbeb3d932e4de66be0c

## Contributed paper

**TARGET AUDIENCE** We target conservation scientists and decision makers addressing Environmental Impact and wildlife policy.

## Supplementary Material

Supplementary material is available in this OSF repository: https://osf.io/r9mkp/?view_only=fb71321265d14dbeb3d932e4de66be0c

## ACKNOWLEDGEMENTS

We thank J. Girard, S. McFarlane, R. Bhatti, S. Hayne, M. Hornseth, and G.D. Sutherland for discussion and suggestions that helped improve this manuscript. We also thank E. McIntire, T. Micheletti, A. Chubaty, and the Predictive Ecology group for their support and contributions to R package development.

